# A vertical stacking approach to rapid generation cycling for indoor growth of tall annual crops - the case of *Vicia faba*

**DOI:** 10.1101/2025.02.09.637347

**Authors:** Maria Pazos-Navarro, Richard G Bennett, Christine Munday, Robert Creasy, Samuel C Catt, Judith Lichtenzveig, Federico Ribalta, Janine Croser

## Abstract

Accelerating breeding cycles through rapid generation turnover is a critical strategy to improve genetic gain in crops. This study presents a vertically stacked accelerated single-seed descent (aSSD) protocol for faba bean (*Vicia faba* L.), aimed at reducing generation time while ensuring robust plant development and seed viability. Alongside photoperiod extension to truncate time to flowering, we present innovations including flurprimidol application to control plant height and enable vertically stacked growth, and optimisation of immature seed desiccation for precocious seed germination. Under controlled environment conditions, the protocol achieved 4.1 to 5.7 generations per year across phenologically diverse germplasm. Plant height was successfully reduced by up to 50% with flurprimidol without compromising flowering or seed yield. The methodology was validated within the University of Adelaide led Australian PBA faba bean breeding program (FBA), advancing 14 recombinant inbred line populations over three years. Beyond faba bean, this scalable, genotype-independent platform offers potential applications for other tall annual species in breeding and vertical farming systems, enabling year-round crop production and genetic improvement. Moreover, the protocol’s adaptability to closed-environment agriculture presents unique opportunities for plant-based life-support systems in extreme environments, including space missions. This integrated approach to speed breeding and vertical farming represents a significant advancement in agricultural innovation, with implications for food security and space exploration.

## Background

The global population is expected to reach 9.7 billion by 2050 (1), posing a challenge to agricultural producers, plant breeders and the research community to accelerate genetic improvement for productivity-related traits in staple crops. The generation length was recognised long ago by animal breeders for its impact on selection effectiveness and, by extension, its effect on genetic improvement rate (2) (3). These observations have equal relevance to plant breeding and impact all parameters of the equation proposed by Fisher (4) whereby the rate of genetic gain (<u>Δ</u>) is a function of selection intensity (*i*), heritability (*h^2^*) and selection differential (*S*). Breeding programs that integrate protocols to speed generation turnover can select and cross more individuals with desirable traits (which impacts on *i*), rapidly identify and propagate individuals with a high heritability for desirable traits or identify and select individuals with superior genetic traits (which impacts on *S*).

As a result, considerable attention has been focused on the development of methods to accelerate the generation cycle across a range of economically important plant species. In 1964, Guha and Maheshwari (5)presented the concept of laboratory-based regeneration from male gametes which led to the development of double haploidy as a technique for fixation of alleles in a single generation in responsive species (as reviewed by (6)). Soon after, Brim (7), building on the ideas of Goulden (8), presented a detailed single seed descent (SSD) method focused on the growth of plants under conditions geared to achieve rapid progression to reproduction rather than vegetative biomass, highlighting its value in rapid fixation of alleles. Since the early 2000’s researchers, spurred on by technological advances in controlled environment plant growth, have further refined SSD by manipulation of photoperiod, temperature, light spectrum, CO_2_ levels, and hydroponic nutrient delivery. These long-standing methodologies recently grouped under the heading of ‘speed breeding’ (9), continued to be applied across public and private breeding programs for a range of broadacre crop species, including cereals, oilseeds, maize and pulses (reviewed by (10)).

For the temperate pulses chickpea (*Cicer arietinum* L.), field pea (*Pisum sativum* L.), lentil (*Lens culinaris* ssp. culinaris), and faba bean (*Vicia faba* L.), which remain recalcitrant to the robust application of doubled haploid techniques, our group has developed and delivered an accelerated Single Seed Descent (aSSD) platform to cycle >25,000 individuals/year across the four species for the Australian national breeding programs (modified from (11) (12) (13)). We have demonstrated the value of combining aSSD with marker assisted selection and advanced field trial analytics to compress chickpea breeding pipelines by up to 3.5 years (13), with phenotypic selection (14) and to fast-track the development of chickpea, lentil and lupin (*Lupinus angustifolius* L.) populations for gene-trait discovery (15) (16) (17) (18). A pipeline of aSSD-derived temperate pulse lines is embedded in the four major pulse improvement programs in Australia and under evaluation for traits of interest.

Vertical stacking forms an integral part of our pulse aSSD platform. The improvements in the design and cost of waterproof, low-heat, light emitting diode (LED) lighting systems that facilitated accelerated generation cycling also provided the opportunity to grow plants in vertically stacked layers under controlled environment conditions. Vertical stacking or vertical farming has attracted attention for plant-based urban food production due to low ‘food miles’, high plant density/m^2^ and potential for application of circularity principles including energy (heat) redistribution, production of biogas from organic waste, water, and nutrient recycling (reviewed by (19)). Closed environment-agriculture will be required to support future off-planet exploration and colonisation and is the subject of current collaborative research between NASA and partners (Plants for Space ARC Centre for excellence, https://plants4space.com) (20). Within research and breeding organisations, vertical stacking of plant growth systems can improve the efficiency of utilisation on a per m^2^ basis of controlled environment space, resulting in substantial running cost reductions and potential for higher plant throughput. To successfully adapt our aSSD protocol to a vertically stacked plant growth framework, we needed to develop methods to manage plant height (21) (22) (23) and achieve reliable flowering and seed set under fully artificially lit conditions.

Here we describe i) a robust aSSD protocol for faba bean integrating *in vitro* or *in vivo* immature seed germination protocols and ii) two methodology refinements aimed at integrating vertical stacking approaches within crop speed breeding-related platforms. The first of this methodology refinements is the control of plant height through a combination of root restriction and exogenous application of anti-gibberellin, with no negative effect on flowering time and seed viability. The second is fully artificially photoperiod supplied by LED arrays with a low red to far red ratio (R:FR) spectrum to improve synchronisation of time to flowering across breeding populations derived from phenologically diverse parents. Faba bean provides an excellent case study for vertical stacked speed breeding due to a field height of up to 1.8 m, a high flower: seed ratio under both field and controlled growing conditions and an outcrossing rate of up to 25-30% (24).

## Material and Methods

All experiments were undertaken at the University of Western Australia, Perth (lat: 31°58′49″S; long: 115°49′7″E). Plants were grown in steam-pasteurised potting mix (pH: 5.8-6.5, UWA Plant Bio Mix - Richgro Garden Products, Australia Pty Ltd – Supplementary Table 2), hand watered daily and fertilised weekly with 20 mL (0.4 L pot) or 30 mL (1 L pot) of a water-soluble N:P:K fertiliser (19:8.3:15.8) with micronutrients (Poly-feed, Greenhouse Grade, Haifa Chemicals Ltd.) at a rate of 16 g L^-1^ (w: v). Each pot contained a single plant and was considered an experimental unit. The plants’ reproductive stage was recorded either at anthesis (when petals are just longer than the sepals), or as open flower. For light quality and quantity measurements we used a Sekonic C7000 SpectroMaster spectrometer (Sekonic Corp., Tokyo, Japan) as per (11). Light intensity values were averaged over three measurements in the range of 400–780 nm: blue (B, 400–500 nm), red (R, 600–700 nm) and far-red (FR, 700–780). Relative (%) and absolute (μmol m^−2^ s^−1^) intensity values in the ranges of B, R and FR were calculated (25). Red to far-red ratio (R:FR) calculations followed the method by (26): photon irradiance between 655-665 nm/photon irradiance between 725-735 nm (Supplementary material: Spectral models and measurements and Supplementary Table 1). All seed was kindly provided by FBA.

For the development of this methodology all statistical analysis were performed using RStudio 2022.12.0, and R version 4.2.3. software and graphs were plotted using the ggplot2 library.

### Effect of antigibberellin on plant height

Field grown faba bean reach a height of up to 1.8 m at full maturity. To facilitate growth in vertically stacked shelves, we tested exogenous application of the antigibberellin flurprimidol (Topflor, SePRO Corporation, USA) to restrict plant height as per (21) (22) (23). Seed of Nura, a shorter variety with mid-maturity, and PBA Samira, a tall variety with early to mid-maturity, were sown in 0.4 L pots and plants were grown in a glasshouse set at 24/18 ⁰C with natural photoperiod supplemented and extended to 18 h by light emitting dioide (LED) arrays AP67 L-series, Valoya, Finland. When plants had reached the 3 - 5 fully reflexed leaf stage, *c.* 15 days after sowing (DAS) (Figure 1A), they were treated with 5 ml flurprimidol in either 0.00, 0.05, 0.10, 0.20, 0.40 or 0.80 % v: v, applied as leaf and soil drench (Figure 1B).

**Figure 1.**
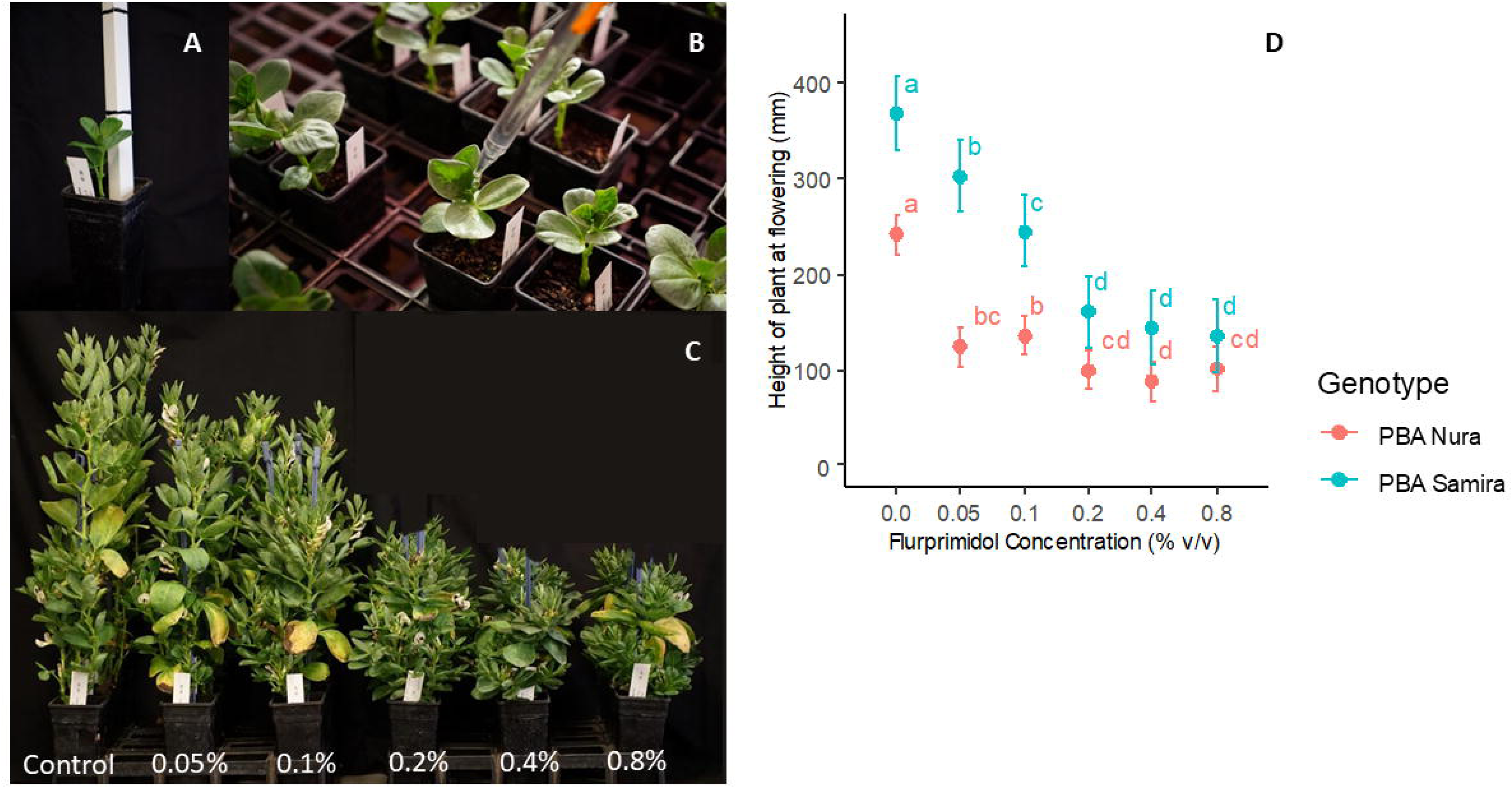
(A) PBA Samira plant at 15 days after sowing (3 fully reflexed leaf stage). (B) leaf and soil drench. (C) Effect of flurprimidol concentration applied to PBA Samira. (D) Effect of flurprimidol concentration applied to Faba bean genotypes PBA Nura and PBA Samira on plant height at flowering (mm). Letters indicate significance groupings (P=0.05) among flurprimidol concentrations within each genotype, and do not compare differences between genotypes.

Four experimental units per line (Nura and PBA Samira) and flurprimidol dose were set in a complete random design (n = 24). The effect of the antigibberellin on the response of plant height at flowering, days to flower, number of seeds, and seed weight were analysed by treating the two genotypes separately and conducting a one-way ANOVA and LSD test (α= 0.05, Agricolae library).

### Effect of photoperiod on floral phenology and seed set

The effect of photoperiod on days to flower (DTF) and seed number per plant was assessed in an early field flowering Australian breeding line ‘1952-1’ and a late variety ‘Icarus’ (93 and 105 DTF respectively when grown in season, in-field at Turretfield Research Centre, South Australia). Seeds were sown in 0.4 L pots and were placed in controlled environment rooms (CERs) (Conviron, model BDW40, Controlled Environments Ltd, Canada) with light intensity set at 500 μmol m^−2^ s^−1^ provided by metal halide lamps (Venture lamp type HIE 400W) and incandescent bulbs, with a R:FR light ratio of 1.3. The day/night temperature of 24/18 ⁰C (12 h: 12 h) was uncoupled from the photoperiod length and remained constant in all seven photoperiod treatments. Due to CER availability, the experiment was performed in two sets. Set 1 = photoperiods 9, 12, 14, 16, 20 h; Set 2 = photoperiods 16, 18, 20, 24 h. Days to flower and number of seeds were recorded per plant. The experimental units (pots) were set in a split-plot design with the photoperiod treatments as main plots with partial repeats (16 and 20 h) and plant lines as secondary plots, n = 3 – 9 randomly distributed per room.

The effect of photoperiod on number of seeds and days to flower in the two genotypes were analysed separately using a one-way ANOVA. Means of response and 95% confidence intervals were calculated for the photoperiod treatments and compared (α= 0.05) using an LSD test (Agricolae library).

### Effect of light spectrum and vertical farming conditions on floral phenology and seed set

To determine the effect of light spectrum on time to flower and number of seed produced per plant. We grew 1952-1 and Icarus, under six different controlled growth environments (E1 – E6) set at constant photoperiod (18 h) and temperature (22/18 ⁰C day: night), but different in light spectrum and intensity (Figure 2). All plants were drenched with 5 ml 0.2% v: v flurprimidol at 15 DAS.

**Figure 2.**
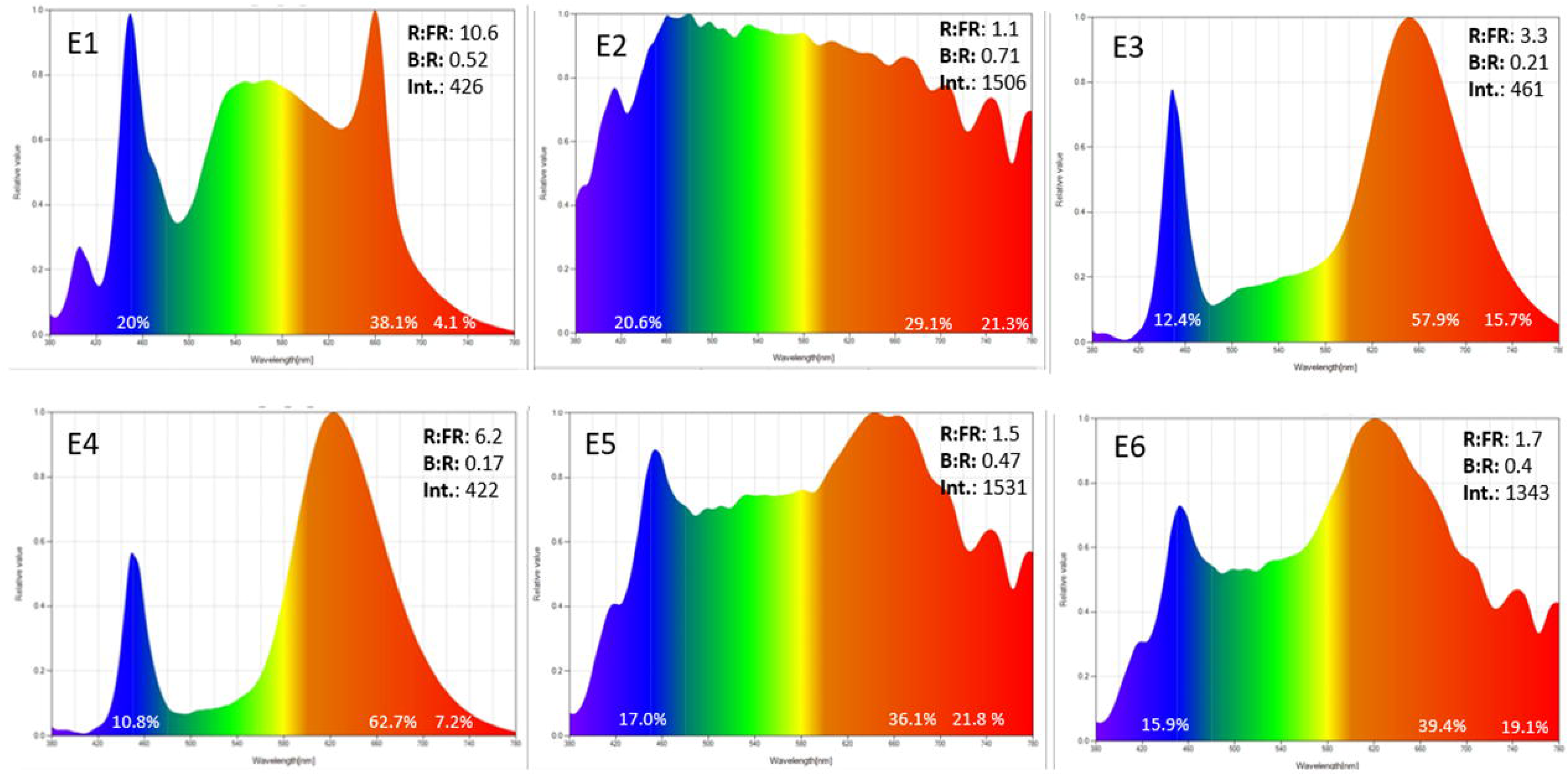
Light spectrum profiles across the six environments used in this study (E1-6). The X axis is wavelength (nm) from 300 to 780 nm, while the Y axis is the strength of that portion of the wavelength relative to the strongest wavelength (0-1). Percentage numbers on each graph represent amount of blue (B), red (R) and far red (FR) nm from the total in each profile, and B: R and R: FR the ratios among those. Int represents the total intensity (photon flux density, μmol m^−2^ s^−1^) in each environment. E1: natural sunlight spectrum NS1 (Valoya RX series); E2: natural sunlight; E3: far-red enriched AP67 spectrum, E4: AP673 spectrum; E5: natural + AP67; E6: natural + AP673. Broad band red (R, 600–700 nm) and far-red (FR, 700– 780 nm) wavelengths with % of total light spectrum values for each region indicated within each box. Narrow band R (650 ± 5 nm) and FR (730 ± 5 nm) regions were used for calculating the ratio of R: FR light.

Light spectrum, intensity, photoperiod, and quantum were central to the analysis of our results. In fully artificial environments E1, E3 and E4, the light features remain constant over the course of the experiments. However, in E2, daylength and therefore light quantum changed over the course of the experiment. In the supplemented and extended photoperiod environments E5 and E6, changes in daylength altered both the quantum and quality of light as mixtures of natural and artificial light varied across the day. Models of the daylength, photoperiod, intensity and light spectrum measures were generated and adapted to each individual’s planting date and date of anthesis, using measurements of light quality and intensity, and daylength during the plant’s growing period as inputs (full details of spectral models and measurements are presented in supplementary material). We tested growth under vertically stacked plant growth conditions in E3 and E4, with pots placed on a three-tiered shelving unit with 80 cm high spacing between shelves and the Valoya AP67 L-series arrays linked to form two parallel lighting arrays, covering a surface of 200 cm x 60 cm per shelf (Figure 3 A and 3B).

**Figure 3.**
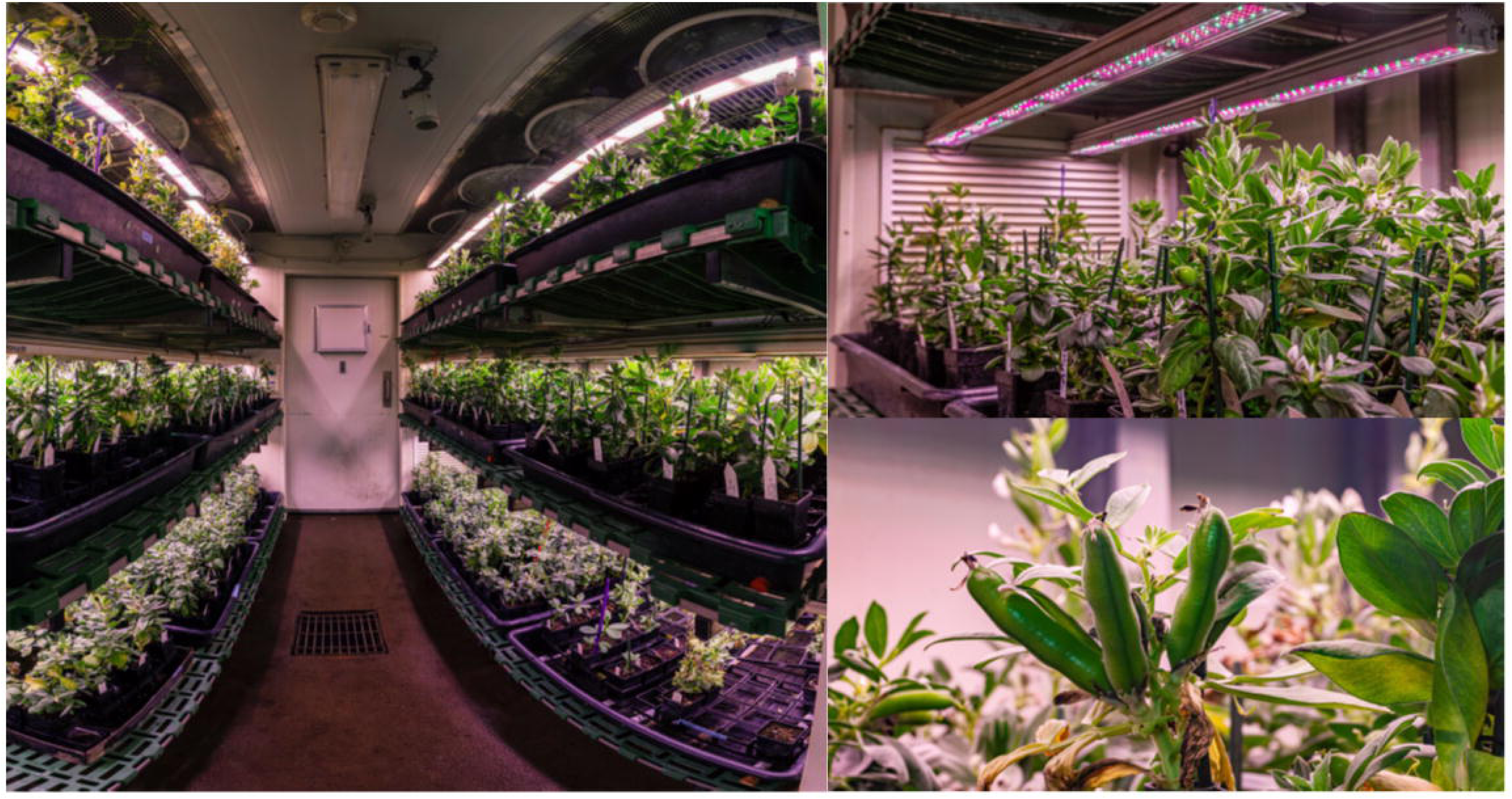
Industry relevant platform for Faba bean. Plants grow in A. multi-tiered shelves with 80 cm high spacing between shelves and B. the Valoya AP67 L-series arrays linked to form two parallel lighting arrays, covering a surface of 200 cm x 60 cm per shelf. C. Detail of Faba bean pods grown under aSSD conditions.

The experimental units (pots) were set in a split-plot design with the environment (E1 – 6) treatments as main plots and plant lines as secondary plots, n = 5 randomly distributed per room. The experiment was repeated twice. Time to flower, number of seed per plant and 100 seed weight were recorded for each line in each environment.

To understand the effect of both pot size and light spectrum on days to flower, number of seeds produced, and 100 seed weight (g), Icarus and 1952-1 were separately subjected to two-way ANOVA with pot size and environment as the explanatory variables. For number of seeds, where both variables were significant and had significant interactions, comparisons between all variable combinations were analysed using one-way ANOVA and LSD test (α=0.05) and plotted as previously described. For days to flower and 100 seed weight, there was no significant pot effect or interaction between environment and pot size, hence, the effects of light environment were analysed using one-way ANOVA, and LSD test (α=0.05). In both cases, only the results for the 0.4 L pots are plotted as this pot size allowed for maximum planting density and, therefore, was adopted in our standard procedure.

In those environments with extended photoperiod, further analyses were conducted of the effect of light spectrum and quantity (intensity) on days to flower, seed number per plant and 100 seed weight by calculating and plotting correlations between these plant growth variables and selected aspects of light quality using ‘cor’ and ‘corrplot’ in R. For this analysis, genotypes were analysed separately, and only those treatments grown in 0.4 L pots were included.

### *In vitro* precocious seed germination

Forty plants of 1952-1 were grown in 0.4 L pots under our accelerated single seed descent (aSSD) conditions of 22/18 ⁰C (coupled to light/night cycles), and 18 h photoperiod provided by Valoya AP67 LED arrays with 0.2% v: v flurprimidol treatment applied at 15 DAS. At anthesis, flowers were tagged at the base of the floral peduncle (APLI Strung Tickets, Pelikan). To determine the association effect between seed developmental stage and robust germination, pods with immature seeds were harvested 26, 28, 30, 31 and 32 days after anthesis (DAA) and seeds were cultured *in vitro* (modified from (4)). Briefly, pods were surface sterilized in 70% ethanol for 1 min, followed by 5 min in sodium hypochlorite (21 g/L active HOCl), and rinsed thrice in sterile water. Pods were opened with a scalpel and forceps under sterile conditions and immature seeds with removed integuments were cultured in 20 x 90 mm petri dishes containing 20 mL of Gamborg B5 basal medium (G398, Phytochemlab) with 20% sucrose, pH adjusted at 5.7 and solidified with 0.6% Type M agar (Sigma). Seeds were considered germinated when both radicle and shoot had emerged. After *ex vitro* transfer, seedlings were rinsed and transferred directly to soil in 0.4 L pots under aSSD conditions as above.

The proportion of germinated seeds 5 days after establishment *in vitro*, seedlings with normal development after 21 d, and plants that reach reproductive stage, as well as days to flower and generation time, were recorded for each treatment (DAA). The experiment was repeated three times with a minimum of 27 seeds per treatment. The experimental design was completely randomized, and the statistical analysis performed using χ2 test for homogeneity of the binomial distribution. A Proportion test analysis was performed when significant differences between treatments were observed (P=0.05).

### Fully *in planta* accelerated SSD platform

For users without access to sterile culture facilities, we developed a fully *ex vitro* aSSD protocol, validated across 30 phenologically diverse individual lines (Supplementary Table 3), representing the variability within the FBA (Pers. comm. Dr Jeff Paull, FBA primary breeder (2007-2021)). In brief, two seeds per line at generation 1 were sown in 0.4 L pots at 22/18 °C (light/dark cycle), with an 18 h photoperiod provided by AP67 Valoya L-series and light intensity of 350-400 μmol m^-2^ s^-1^ as per (11). After seedling establishment and prior to antigibberellin treatment, the number of seedlings were reduced to one. Flurprimidol at 0.2% v: v was applied as a 5 mL leaf drench at 15 DAS. Flowers were tagged at anthesis, and immature seeds harvested at physiological maturity, c. 32 days after anthesis. The in-pod immature seeds were dried in seed envelopes placed in seedling trays in a *c*. 3 cm deep bed of orange indicator silica gel (Dessico Pty Ltd, Australia) for 5-7 days at 25 °C (Thermoline cabinet). At 8% seed moisture, measured with an active water meter (Rotronics HC2-AW-USB, www.rotronic.com) and converted to seed moisture reading (as per (27)), seed were resown in pots and grown under the same conditions as the previous generation. The seeds from the final generation were left an additional 14 -21 days to mature naturally on the plant. From each individual line one pot was sown, except for breeding line 1952-1 (early flowering), PBA Warda (mid flowering) and Icarus (late flowering). For these lines, a minimum of 30 plants were grown to establish if there was variability within the lines used. From each line, days to flower and time to harvest were recorded, and generation turnover time estimated.

### Methodology validation within FBA

Over three years, the aSSD platform progressed (F_2:5_) 14 faba bean recombinant F_2_ populations segregating for important agronomic traits from the FBA, for which the pedigree was recorded for each recombinant inbred line (RILs) and spare seeds were made available to the breeders for every line and generation. Plants were subjected to the same growth conditions as described in “Fully *in planta* accelerated SSD platform’. For each individual in each generation, flowering time was recorded, and pods removed 32 DAA. Seeds were desiccated in pods as described previously, and three seeds from each RIL were resown and thinned randomly to one individual per pot post seedling establishment.

In the example provided here, we selected for herbicide resistance at F_4_ generation. The selection for herbicide resistance was performed on 21-day old F_4_ seedlings. Plants were treated with the herbicide metribuzin (180 g active ingredient ha^-1^) and then leaf damage was scored after a further 14 days to select tolerant RILs which were then progressed for F_5_ seed collection.

## Results

### Effect of photoperiod and genotype on reproductive phenology under controlled-environment conditions

Faba bean is a photoperiod responsive species. Under controlled-environment conditions of photoperiod and temperature, accessions which are considered ‘early’ and ‘late’ under field conditions differ in their reproductive phenology response to photoperiod. The breeding line 1952-1 (‘early’) showed a typical facultative long-day response as it successfully flowered under short days and time to flower was shorter in longer photoperiods (Figure 4A). The cultivar Icarus had an obligatory long day response, it failed to flower at short photoperiods (9 and 12h), and its days to flower (DTF) shortened exponentially with longer photoperiod treatments. The accessions reached photoperiod saturation at different treatments, 1952-1 at 16 h, and Icarus at 20 h. Neither of the lines set seed under short days (9 h; Figure 4B). We selected an 18 h photoperiod for the aSSD platform as it resulted in acceleration of flowering while maintaining adequate seed production for SSD purposes and general plant health.

**Figure 4.**
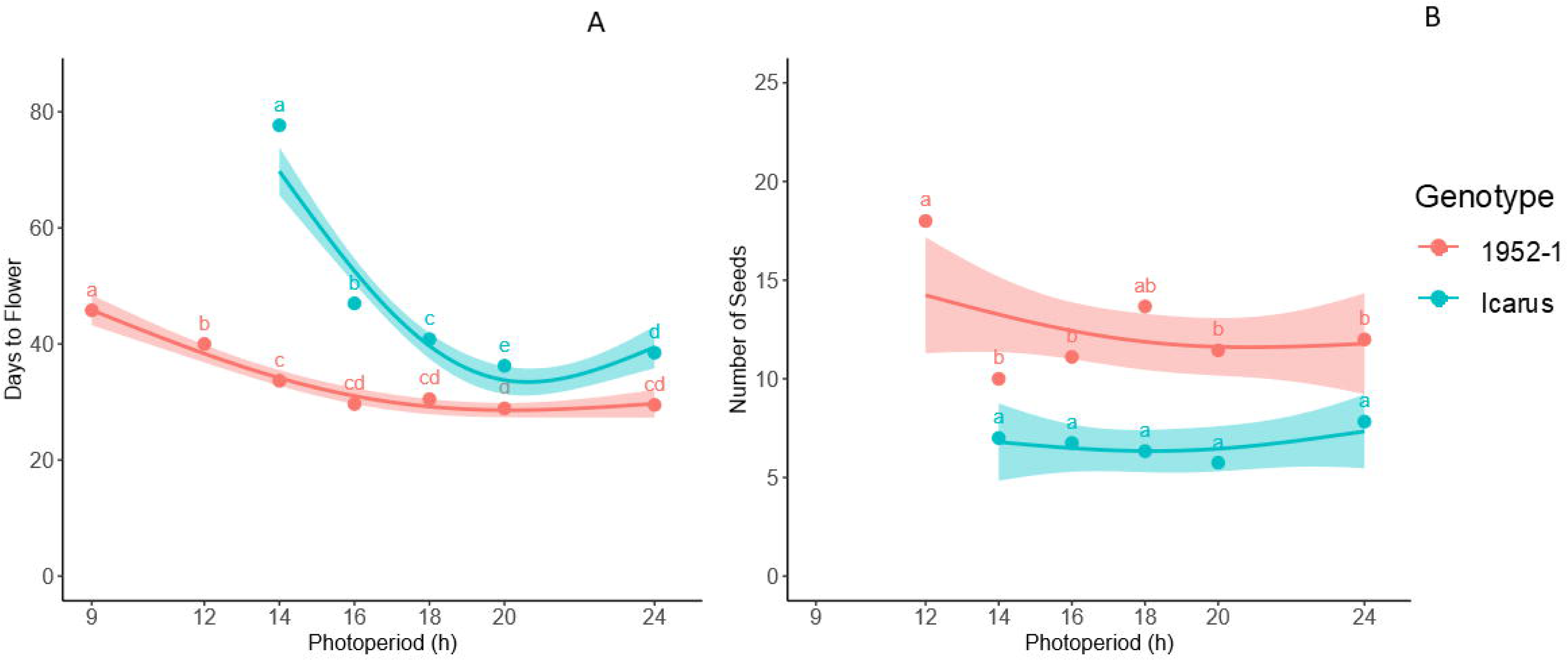
Effect of photoperiod on (A) days to flower and (B) number of seeds produced per plant. Letters indicate significance groupings (α= 0.05) among photoperiods within each genotype, and do not compare differences between genotypes.

### Plant miniaturisation is achieved using a single antigibberellin treatment

Decreased plant height during plant development was achieved with increasing flurprimidol concentrations (P<0.001) (Figure 1C and 1D), in both naturally tall (PBA Samira) and short (Nura) plants. The 0.2% v: v concentration applied as 5 mL leaf drench at 15 DAS was selected due to its ‘goldilocks’ miniaturisation effect, reducing height of PBA Samira plants from *c*. 380 mm to *c*. 180 mm, and with no significant effect on time to flower (P>0.05), number of seed produced per plant (P>0.48) or seed weight (P>0.10) (Supplementary Figure 1A, 1B and 1C). The 0.2% v:v concentration and timing of flurprimidol application was incorporated as a routine step in the aSSD protocol. The resulting miniature plants allowed for growth in 0.4 L pots and a density of up to 80 plants/m^2^ in all environments, including the vertically stacked shelves (Figure 3).

### Minimal effect of light spectrum at photoperiod saturation

The photoperiod effect was more pronounced for Icarus with an average of 27 d truncation in time to flowering in the extended photoperiod environments (a 39% reduction compared to natural photoperiod in E2), while 1952-1 was 7 d shorter (21 % reduction) (Figure 5A).

**Figure 5.**
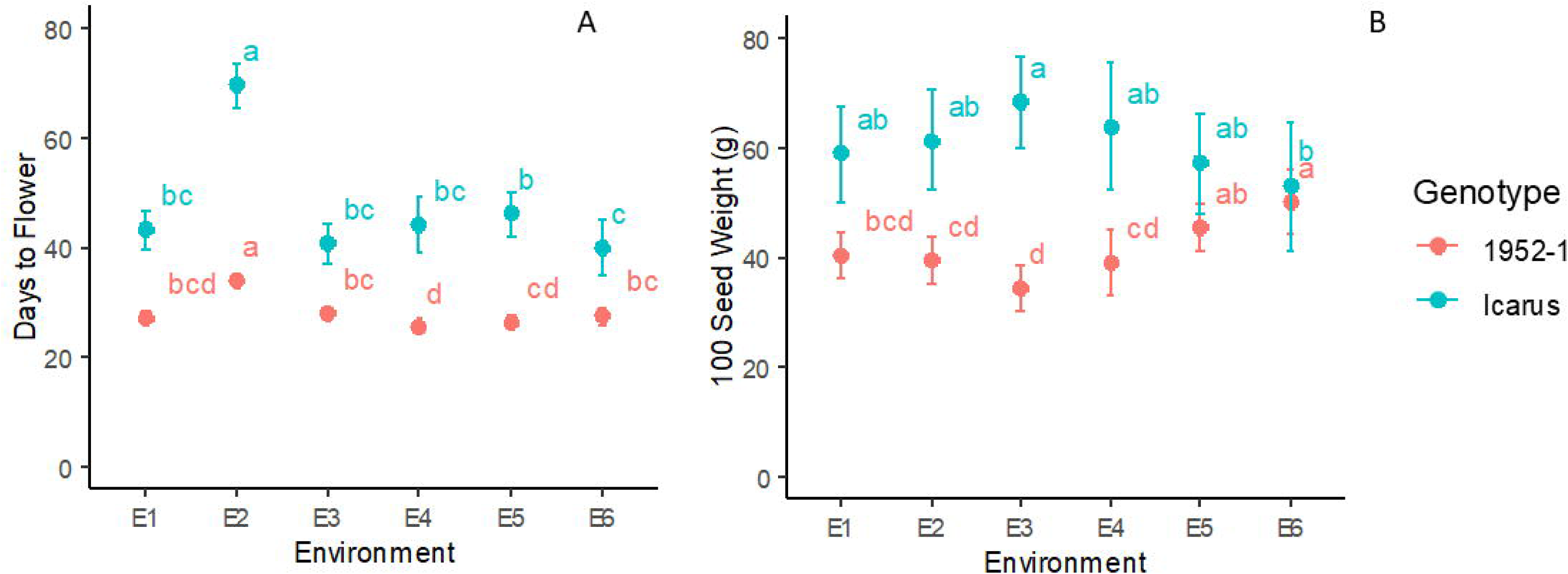
Effect of light environment (E1: natural sunlight spectrum NS1 (Valoya RX series); E2: natural sunlight; E3: AP67 spectrum, E4: AP673 spectrum; E5: natural + AP67; E6: natural + AP673) on (A) days to flower and (B) 100 seed weight (g) in Faba bean genotypes Icarus and 1952-1. Letters indicate significance groupings (α= 0.05) among environments within each genotype, and do not compare differences between genotypes.

Within fully artificial (E1, E3 and E4) or semi-artificial light environments (E5 and E6), spectrum had a bearing on days to flower, however the effects were relatively small. E3 resulted in a longer time to flower than E4 for 1952-1, while for Icarus E5 was a longer time than E6 (Figure 5A and Supplementary Table 3). For 1952-1, the light spectra had a substantial bearing on 100 seed weight, with the heaviest seeds produced in semi-artificial lit environments (E5 and E6). For Icarus, no clear effects were observed (Figure 5B and Supplementary Table 3).

Pot size is an important factor in determining the plant density within controlled environments. Growth in 0.4 L versus 1 L pots did not change days to flower in 1952-1 or Icarus (Table 1). Unsurprisingly, the larger pot size (1L) led to increased seed number per plant across most environments (Figure 6A and 6B and Supplementary Table 3). Notably, the pot size effect was smallest in E2 (the natural light). Overall effects of pot size on seed weight were negligible (Supplementary Table 3). We regularly use 0.4 L pots, as SSD requires only one viable seed to move to the next generation, and it results in the production of 2-5 spare seeds as backup in case of germination failure in subsequent generations.

**Figure 6.**
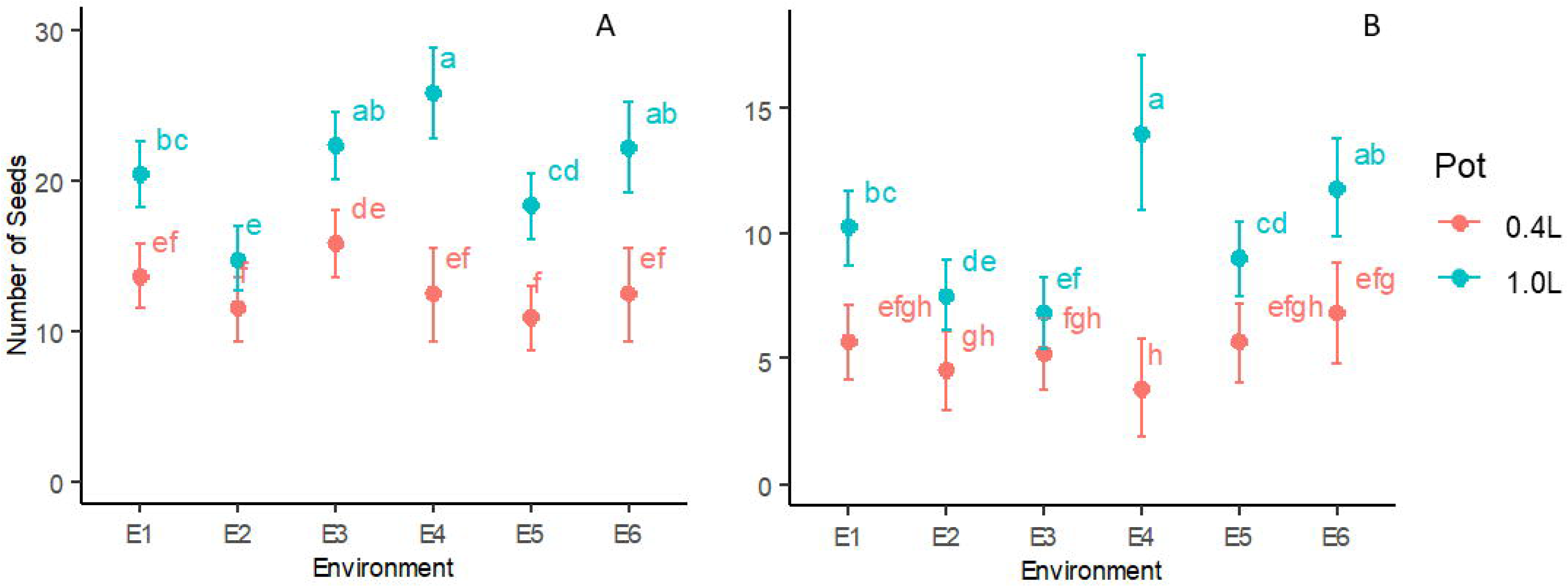
Effect of light environment (E1: natural sunlight spectrum NS1 (Valoya RX series); E2: natural sunlight; E3: AP67 spectrum, E4: AP673 spectrum; E5: natural + AP67; E6: natural + AP673) and pot size on the count of seed produced per plant by faba bean genotypes (A) 1952-1 and (B) Icarus. Letters indicate significance groupings (P=0.05) among all environment and pot size combinations within each genotype.

**Table 1.** Effect of seed developmental stage (26-32 days after anthesis, DAA) on the proportion of seeds with normal seedling development 21 days after *in vitro* culture, and proportion of plants from the total number of cultured seeds reaching reproductive stage up to day 50. Statistical analysis was performed using χ^2^ test for homogeneity of the binomial distribution (χ^2^= 10.85, df = 4, P = 0.03) and Prop.Test, different letters represent significant differences between seed developmental stage (DAA) (P ≤0.05; n> 27).

| Seed developmental stage (DAA) | Proportion of seeds that developed into seedlings | Proportion plants that reached floral stage |
| --- | --- | --- |
| 26 | 0.67 b | 0.67 b |
| 28 | 0.90 a | 0.60 b |
| 30 | 1 a | 0.73 ab |
| 31 | 1 a | 0.84 ab |
| 32 | 0.93 a | 0.93 a |

### Truncation of seed development without affecting germination and plant development

The age at which the immature seeds were harvested did not affect gemination, as root and shoot growth was observed in all seeds harvested independently of their DAA stage (Supplementary Figure 2A and 2B). However, the proportion with normal seedling development and proportion of reaching flowering depended on the DAA (Table 1). Reliable seedling establishment was reliably achieved from plants derived from immature seeds older than 28 DAA (over 90%), while subsequent flowering was achieved from 32 DAA seeds (Table 1).

In a similar manner, Generation 1 time (i.e., G1 - seed sowing to immature seed culture) and subsequent generations (G2+, immature seed culture to immature seed harvest of the following generation) were affected by the immature seed developmental stage (Table 2, P = 0.001). While the culture of immature seeds 26 DAA achieved the shortest G1 and G2+ generation cycles, it also yielded the lowest proportion of seeds reaching the reproductive stage. Delaying the harvest of immature seeds until 32 DAA resulted in more robust precocious seed development and subsequent flowering and enabled G1 and G2+ generation cycles below 70 days. Supplementary Figure 2 shows a snapshot of the *in vitro* process from precocious seed gemination to seed harvest of the following generation.

**Table 2.** Plant cycle length for the first generation (G1, mature seed sowing to immature seed culture) and subsequent generations (G2+, immature seed culture to immature seed harvest of the following generation) following *in vitro* culture of immature seeds at embryo physiological maturity (26-32 days after anthesis, DAA). Different letters represent significant differences between DAA.

| Seed developmental stage (DAA) | G1 (days) | G2+ (days) |
| --- | --- | --- |
| 26 | 63.1 d | 67.8 b |
| 28 | 64.4 c | 71.2 a |
| 30 | 66.0 b | 71.0 a |
| 31 | 68.2 a | 71.7 a |
| 32 | 68.1 a | 69.6 ab |

### Protocol validation with diverse germplasm

We validated our fully *in vivo* aSSD protocol with a range of phenologically diverse germplasm, selected by the breeder to represent the genetic variability within the FBA program (Supplementary Table 4) using the above optimised conditions. Under aSSD conditions the different lines flowered 25 to 51 days after sowing, and the variability within a line based on 1952-1 (early flowering genotype), was 3 days for Warda (mid flowering) and 6 for Icarus (late genotype).

In our *in vivo* protocol we established the overall generation time as the summation of days to flower, days to harvest of the immature seed (32 days) and desiccation treatment (5 - 7 days). Across the diverse germplasm evaluated, the proposed protocol resulted in generation cycles between 64-90 days, enabling the progression of 4 – 5.7 generations per year (Supplementary Table 4).

We translated the research into an industry relevant platform (Figure 3) which was validated utilising 14 F_2_ populations provided by the FBA over the years 2018 – 2021As a practical case study of the application of this platform we hereby dissect the time it took to progress from F_2_ to F_5_ for a metribuzin tolerance population, with plants subjected to metribuzin tolerance selection in the F_4_ generation. In brief, 73 F_2_ independent lines were sown, and aSSD used to cycle to F_4_. In the F_4_ generation, 21-day-old seedlings were sprayed with metribuzin, and leaf damage was evaluated 14 days post spraying. The surviving plants were cycled using aSSD to produce F_5_ seed. Twenty-eight elite RILs were returned to FBA for further selection. These three generation cycles including selection were achieved in 342 d. It is worthy of note that this example population was slower to progress through the generations, as we allowed extra time for F3 plant development and F4 seed maturation. This ensured the health of F4 seed to enable accurate screening for herbicide tolerance.

## Discussion

We have demonstrated the effective integration of vertical farming and speed breeding to accelerate generation cycle turnover in the globally significant, protein rich pulse crop faba bean. This approach can improve the rate of genetic gain for crop species and maximise research space utilisation in costly controlled environments. Combining vertically stacked growth with antigiberellin-mediated plant height restriction enables simultaneous growth of large breeding populations, increasing the probability of identifying desirable recombinants. A vertically stacked approach to speed breeding also offers opportunity for direct technology transfer from advances made in vertical farming including robotics for tasks such as planting, watering, fertilizing, monitoring, and harvesting.

Temperature along with duration and intensity of supplied photoperiod are the undisputed key factors in successful generation cycling. Temperature, light, and water/nutrient availability interact with plant genetic pathways to determine the timing of flowering. Speed breeding protocols rely on triggering adaptive plasticity in response to changes in environmental cues, particularly light. This is typically achieved through saturation of photoperiod, which can be easily achieved by supplementing natural light in a glasshouse with conventional (metal halide/fluorescent/incandescent/high pressure sodium) or newer (light emitting diode, LED) lighting technology. In faba, we identified variation in photoperiod saturation between the early (16 h) and late (18 h) field flowering genotypes. For the early type, 1952-1, 9 h photoperiod was sufficient to trigger flowering, but no seed set; and 12 h resulted in the highest seed number, likely due to additional time in the vegetative phase to develop the biomass to support reproduction.

Advances in LED systems enable wavelength customisation and a more nuanced assessment of the role of light quality in speed breeding platforms. In temperate pulses, we have previously identified acceleration of flowering in response to enrichment in the far-red spectrum (11) (12), which we postulate is the result of triggering aspects of ‘shade avoidance syndrome’ (28). For faba bean, we demonstrate here successful plant growth under fully artificial light environments across a wide range of R:FR ratios (viz. 10.2; 3.3; 1.2). We conclude that faba appears relatively insensitive to light quality at photoperiod saturation. Further study of light quality at 12 h photoperiod would likely facilitate better discrimination between R:FR regimes. However, for the purpose of speed breeding within a vertical stacking framework, AP67 arrays (R:FR 3.3, Valoya, Finland) were well-suited to acceleration of flowering, compression of flowering time phenology differences, adequate seed set quantum and viable seed weight.

Miniaturisation of tall plants such as faba bean is essential to the implementation of vertically stacked plant growth. For faba, we achieved a highly reduced mature plant height by chemical application of flurprimidol at the four-leaf growth stage, combined with growth in small pots of either 1 L or 0.4 L. Ochatt (21) proposed flurprimidol to keep field pea (*Pisum sativum* L.) plants at a manageable size in the glasshouse. We adapted this concept and used flurprimidol as an *in vitro* treatment in our effort to develop a genotype-independent, fully *in vitro* rapid generation turnover system for pea (22). This idea was taken up by Mobini (23) to develop an *in vitro* generation acceleration system for lentil (*Lens culinaris* L.) and faba bean. As outlined herein, flurprimidol, applied at the correct concentration and growth stage, is a reliable method of reducing internode length and thus plant height, without extending time to flowering. For the purposes of SSD which requires only 1 viable seed to move to the next generation and 2-3 seeds as generational back-ups, combining flurprimidol treatment with a 0.4 L pot size provided sufficient seed in both the early and late flowering genotypes with negligible effect on days to flowering and weight per seed.

There are two key points in the plant lifecycle where time can be saved, progression to flowering and seed maturation. Building on our studies in other pulses and legume pastures (11) (29) (12) (18) (13), we describe both an *in vitro* and *in vivo* method for achieving robust precocious germination of immature seed following embryo physiological maturity in faba bean. Both methods are simple and relatively easy to implement at scale. The *in vivo* desiccation method is our favoured approach due to not requiring specialized sterile culture facilities. Drying down the in-pod immature seed to a seed moisture content of ∼7.5% (as per the sorption isotherms (27)) enabled reliable germination of seed harvested pre-maturity and saved 1-2 weeks per generation. We observed no germination penalty between mature and immature seed post treatment. However, if generation turnover time is not critical, it is a simple process to stop watering and allow seed to dry down on the plant.

The compression of breeding generation cycle has emerged as a cost-effective tool for manipulating the rate of genetic gain (15) (30). For faba bean, the accelerated single seed descent (aSSD) platform described here has been applied within FBA since 2019. Based on the principles of single seed descent (8) (7), plants are grown under photoperiod extension, late-spring temperature regime and precocious germination technology to enable rapid lifecycle completion year-round for faba, irrespective of the field flowering phenology. Harnessing out-of-season accelerated generation turnover enables i) phenotyping in-season at a more advanced generation and ii) marker-assisted selection (MAS) for rapid identification and culling of lines not homozygous for a desired trait prior to in-season field evaluation. The combined aSSD-MAS approach is a highly efficient method to rapidly introgress novel traits and target downstream breeding program resource allocation.

## Conclusions

The positive implications of rapid generation cycling for rate of genetic gain in crops are widely discussed in the literature (e.g., (31) (30) (32)). Speed breeding (9) has become an international catch phrase to describe a range of techniques including Rapid Generation Cycling (RGC), Fast Generation Cycling (FGC) and Rapid Generation Turnover (RGT). Despite the literature being replete with examples of individual breeding/pre-breeding technologies with potential to this end, there remains no evidence in faba bean of protocols that are i) genotype-independent and ii) delivered at scale across segregating populations in public breeding. We bridge this gap with the development of the accelerated single seed decent (aSSD) platform for faba bean and its extensive delivery to FBA. We provide evidence that faba bean is relatively insensitive to R:FR ratio at photoperiod saturation, negating the need for highly specialised lighting arrays for accelerating generation turnover. We provide a robust protocol for the growth of miniaturised plants to maturity, enabling adoption of vertically stacked plant growth systems that can be adapted to the growth of a range of tall crop plants for plant genetic improvement purposes.

## List of abbreviations

aSSD: accelerated single seed descent
MAS: Marker Assisted Selection
DTF: Days to flower
CER: Controlled environment room
R:FR: red to far red
LED: light emitting diode
DAS: Days after sowing
DAA: Days after anthesis
RIL: Recombinant inbred line
FBA: Australian PBA faba bean breeding program (FBA)

## Declarations

### Ethics approval and consent to participate

Not applicable

## Consent for publication

Not applicable

## Availability of data and materials

The datasets used and analysed during the development of this methodology are available from the corresponding author upon request.

## Authors’ contributions

JC and FR conceived the study. JC supervised the project. FR, JC and JL designed the experiments. The Australian PBA faba bean breeding program (FBA) represented in this manuscript by SC provided all faba bean lines and populations used during the development of this methodology. CM, RB, FR and MPN performed the experiments. RC managed plant growth facilities. RB and FR analysed the data. MPN prepared the manuscript. All authors have edited and participated in the writing process of the manuscript and agreed to participate.

## Acknowledgements

We thank Prof. William Erskine for his leadership, Mr Bill Piasini and Mr Leon Hodgson for glasshouse expertise and Ms Kylie Edwards, Ms Sabrina Tschirren, Ms Karen Nelson, Dr Theo Pfaff-Lichtenzveig and Ms Simone Wells for technical assistance. The authors are indebted to Dr Jeff Paull for providing breeding materials.

