## Supplemental Tables and Figures for "A vertical stacking approach to rapid generation cycling for indoor growth of tall annual crops - the case of *Vicia faba*"

**Supplementary material**

**Spectral models and measurements**

To build models of the light intensity and quality in the natural photoperiod environment and mixed natural and artificial light environments, we used four inputs: a separate single measure of peak irradiance and spectral profile for the natural light and artificial lights; a profile of the change in irradiance of natural light over the course of a day; the daylength of each day during the experiment; and the planting and anthesis dates for the plants grown in the experiments. Light quality and peak irradiance were measured at noon during October in all environments, and after dark in mixed light environments (E5 and E6). The noon measurement in mixed environments was made with the artificial lights switched off. Changes in natural light intensity over the course of a day in daylit environments was measured by logging with Hobo MN2022 light intensity loggers (Onset Computer Corporation, Massachusetts, USA)). Daylength measures were obtained from BOM historical data (www.bom.gov.au.).

Two week's data of logged diurnal intensity measures were averaged on a 5-minute basis to build an average daily profile of light intensity during the daylight hours, and then this model was simplified by splitting the daylit hours into three phases. Dawn and sunrise were captured in the first three daylit hours, sunset and dusk were captured in the last 3 daylit hours, and the remaining daylit hours, the period of which varied with natural daylength, were nominated as a 'full sun' period. The average light intensity for these three periods was compared to the noon measures, to provide a single intensity figure for the dawn intensity, dusk intensity, and full sun intensity. Both the dawn and dusk intensity was found to be 41% of the midday measure, and the intensity of the full sun period was 78% of the midday measure. These ratios were applied to the daylength of each day of the experiment to provide a realistic measure of the total flux received by plants in fully or partially daylit environments for each day between planting and flowering, and to calculate the flux received in the blue, red, and far-red spectra. For mixed environments, these figures were added to the flux provided by artificial lights that ran for 18 hours to provide figures for total intensity and the various measures of light quality.

In order to compare intensity differences among environments, independent of daylength, the intensity of light received during lit hours was averaged over the 24 hour period to provide an average daily intensity figure, i.e. the artificially lit E1 with 18 hours at 426 $\mu$mol $m^{-2}$ $sec^{-1}$ averaged to a daily measure of 319 $\mu$mol $m^{-2}$ $sec^{-1}$, cv. the naturally lit E2 where 12.5h at 1853 $\mu$mol $m^{-2}$ $sec^{-1}$ averaged to 584 $\mu$mol $m^{-2}$ $sec^{-1}$ on a daily basis.

For all light measurements we used a Sekonic C7000 SpectroMaster spectrometer (Sekonic Corp., Tokyo, Japan) as per Croser et al. (2016). Light intensity values were averaged over three measurements in the range of 400–780 nm: blue (B, 400–500 nm), red (R, 600–700 nm) and far-red (FR, 700–780). Relative (%) and absolute ($\mu$mol $m^{-2}$ $s^{-1}$) intensity values in the ranges of B, R and FR were calculated (Cope and Bugbee 2013). Red to far-red ratio (R:FR) calculations followed the method by Runkle and Heins (2001): photon irradiance between 655-665 nm/photon irradiance between 725-735 nm (Supplementary Table 1).

**Supplementary Table 1.** Modelled Light Spectral characteristics of the six environments used in this research. E1: natural sunlight spectrum NS1 (Valoya RX series); E2: natural sunlight; E3: AP67 spectrum, E4: AP673 spectrum; E5: natural + AP67; E6: natural + AP673. Broad band red (R, 600–700 nm) and far-red (FR, 700–780 nm) wavelengths with % of total light spectrum values for each region indicated within each box. Narrow band R (650 ± 5 nm) and FR (730 ± 5 nm) regions were used for calculating the ratio of R:FR light.

| | **Modelled Light Environment** | | | | | | | | | | | |
|---|---|---|---|---|---|---|---|---|---|---|---|---|
| | **Intensity (daily average)** | | **% Blue** | | **% Red** | | **% Far red** | | **B:R wide** | | **R:FR Narrow** | |
| | **Min** | **Max** | **Min** | **Max** | **Min** | **Max** | **Min** | **Max** | **Min** | **Max** | **Min** | **Max** |
| **E1** | 319 | | 0.20 | | 0.38 | | 0.04 | | 0.52 | | 10.62 | |
| **E2** | 584 | 674 | 0.21 | | 0.29 | | 0.21 | | 0.71 | | 1.10 | |
| **E3** | 346 | | 0.12 | | 0.58 | | 0.16 | | 0.21 | | 3.34 | |
| **E4** | 317 | | 0.11 | | 0.63 | | 0.07 | | 0.17 | | 6.16 | |
| **E5** | 706 | 817 | 0.175 | 0.179 | 0.38 | 0.40 | 0.192 | 0.195 | 0.48 | 0.51 | 1.52 | 1.61 |
| **E6** | 715 | 732 | 0.169 | 0.169 | 0.42 | 0.42 | 0.159 | 0.160 | 0.45 | 0.46 | 1.86 | 1.88 |

**Supplementary Table 2.** UWA Plant Bio Mix (Richgro Garden Products) soil composition.

| **Soil Component** | |
|---|---|
| Composted pine bark (fine) | 2.5m3 |
| Coco peat | 1.0m3 |
| River sand quartz | 1.5m3 |
| Limestone (extra fine) | 10kg/ 5m3 |
| Dolomite (Watheroo) | 10kg/ 5m3 |
| Iron Sulphate (Hepta) | 4kg/ 5m3 |
| Iron Chelate | 250g/ 5m3 |
| Osmoform premix (fertiliser) | 2kg/ 5m3 |
| Richgro Gypsum | 1kg/ 5m3 |
| pH: Approx 5.8 - 6.2 | |

**Supplementary Table 3**. Statistics of the effect of growth conditions (environment), pot size and their interactions on time to flower, number of seeds produced per plant and seed weight. The magnitude of effects can be seen in Figures 4 and 5. The effects where P<0.05 are highlighted in bold.

| **Genotype** | **Trait** | **Environment effect (P value)** | **Pot effect (P value)** | **Interaction (P value)** |
|---|---|---|---|---|
| 1952/1 | Time to Flower | **$<2.0 \times 10^{-16}$** | 0.57 | 0.12 |
| | Seed Count per Plant | **$8.8 \times 10^{-07}$** | **$<2.0 \times 10^{-16}$** | **0.0081** |
| | 100 Seed Weight | **$2.68 \times 10^{-05}$** | 0.45 | 0.16 |
| Icarus | Time to Flower | **$<2.0 \times 10^{-16}$** | 0.88 | 0.45 |
| | Seed Count per Plant | **0.002** | **$1.13 \times 10^{-11}$** | **0.0043** |
| | 100 Seed Weight | 0.17 | 0.075 | 0.69 |

**Supplementary Table 4.** Dissection of generation time (including days to flower, days to harvesting and desiccation treatment) across a range of phenologically diverse germplasm, representing the genetic variability within the Australian faba bean breeding program. *Genotype selected for protocol development.

| Genotype | Days to Flower | Days to seed harvesting | Desiccation treatment | Generation time | Generation per year |
|---|---|---|---|---|---|
| **AF08108** | 25 | 32 | 7 | 64 | 5.7 |
| **AF02002-2** | 28 | 32 | 7 | 67 | 5.4 |
| **1714-2** | 29 | 32 | 7 | 68 | 5.4 |
| **AF03001-1** | 30 | 32 | 7 | 69 | 5.3 |
| **DOZA** | 30 | 32 | 7 | 69 | 5.3 |
| **PBA NASMA** | 30 | 32 | 7 | 69 | 5.3 |
| **622-1** | 31 | 32 | 7 | 70 | 5.2 |
| **NURA** | 32 | 32 | 7 | 71 | 5.1 |
| **CAIRO** | 32 | 32 | 7 | 71 | 5.1 |
| **1952-1*** | 33.89 ± 5.88 | 32 | 7 | 72.89 ± 5.88 | 5.0 |
| **PBA WARDA*** | 36.29 ± 5.87 | 32 | 7 | 75.29 ± 5.87 | 4.8 |
| **AF03063-1** | 32 | 32 | 7 | 71 | 5.1 |
| **AF06125** | 32 | 32 | 7 | 71 | 5.1 |
| **AF07125** | 32 | 32 | 7 | 71 | 5.1 |
| **PBA MARNE** | 32 | 32 | 7 | 71 | 5.1 |
| **1322-2** | 33 | 32 | 7 | 72 | 5.1 |
| **AF08207** | 33 | 32 | 7 | 72 | 5.1 |
| **AF08014** | 34 | 32 | 7 | 73 | 5.0 |
| **1727-3** | 35 | 32 | 7 | 74 | 4.9 |
| **PBA SAMIRA** | 35 | 32 | 7 | 74 | 4.9 |
| **FARAH** | 36 | 32 | 7 | 75 | 4.9 |
| **AF05059-1** | 37 | 32 | 7 | 76 | 4.8 |
| **AF04053-1** | 37 | 32 | 7 | 76 | 4.8 |
| **PBA RANA** | 38 | 32 | 7 | 77 | 4.7 |
| **ICARUS*** | 42.47 ± 2.88 | 32 | 7 | 81.47 ± 2.88 | 4.5 |
| **AF06104** | 38 | 32 | 7 | 77 | 4.7 |
| **AF03109-1** | 39 | 32 | 7 | 78 | 4.7 |
| **Kareema** | 41 | 32 | 7 | 80 | 4.6 |
| **1477-4** | 42 | 32 | 7 | 81 | 4.5 |
| **AF05069 TF** | 51 | 32 | 7 | 90 | 4.1 |

**Supplementary Figure 1.** Effect of flurprimidol concentration applied to Faba bean genotypes PBA Nura (short) and PBA Samira (tall) on (A) days to flower, (B) average weight (mg) of 100 seeds and (C) number of seeds per plant. Letters indicate significance in groupings (P= 0.05) among flurprimidol concentrations within each genotype, and do not compare differences between genotypes.

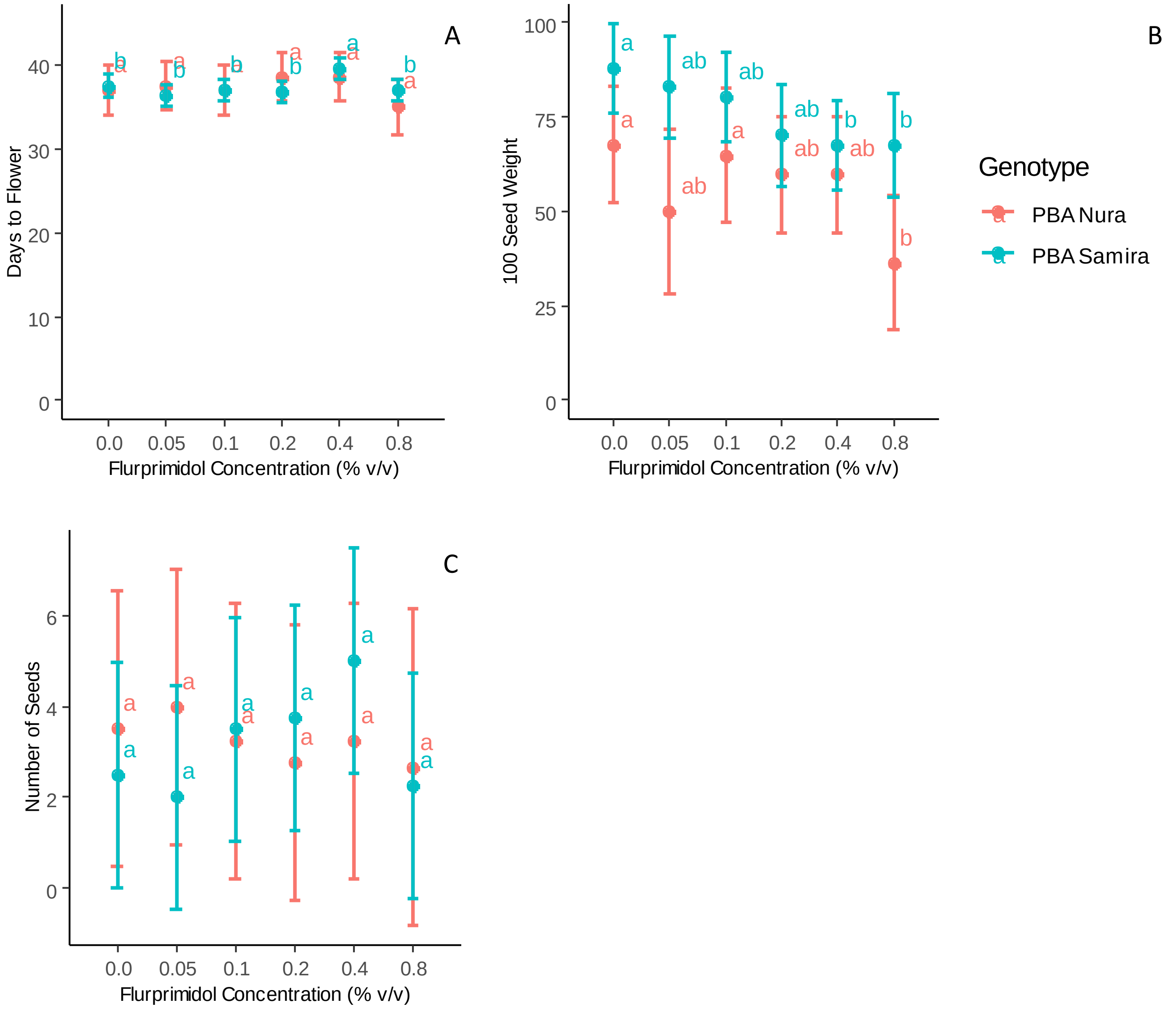

**Supplementary Figure 2**: Immature seed *in vitro* culture of breeding line 1952-1 around embryo physiological maturity. (A) Germinated seeds harvested at 26, 28, 30, 31, 32 days after anthesis (DAA). (B) Geminated seeds *in vitro*. (C) Transfer to *in vivo* conditions. (D) Plants derived from immature seeds flowering.

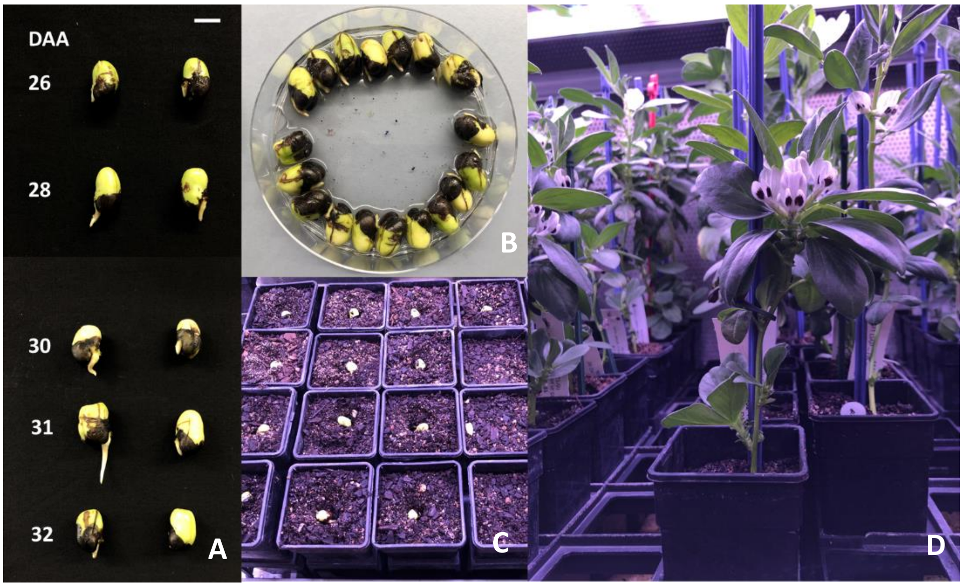
